# Methyltransferase-like 3 modulates severe acute respiratory syndrome coronavirus-2 RNA N6-methyladenosine modification and replication

**DOI:** 10.1101/2020.10.14.338558

**Authors:** Xueyan Zhang, Haojie Hao, Li Ma, Yecheng Zhang, Xiao Hu, Zhen Chen, Di Liu, Jianhui Yuan, Zhangli Hu, Wuxiang Guan

## Abstract

The coronavirus disease 2019 pandemic caused by severe acute respiratory syndrome coronavirus-2 (SARS-CoV-2) is an ongoing global public crisis. Although viral RNA modification has been reported based on the transcriptome architecture, the types and functions of RNA modification are still unknown. In this study, we evaluated the roles of RNA N6-methyladenosine (m^6^A) modification in SARS-CoV-2. Our methylated RNA immunoprecipitation sequencing (MeRIP-Seq) analysis showed that SARS-CoV-2 RNA contained m^6^A modification. Moreover, SARS-CoV-2 infection not only increased the expression of methyltransferase-like 3 (METTL3) but also altered its distribution. Modification of METTL3 expression by short hairpin RNA or plasmid transfection for knockdown or overexpression, respectively, affected viral replication. Furthermore, the viral key protein RdRp interacted with METTL3, and METTL3 was distributed in both the nucleus and cytoplasm in the presence of RdRp. RdRp appeared to modulate the sumoylation and ubiquitination of METTL3 via an unknown mechanism. Taken together, our findings demonstrated that the host m^6^A modification complex interacted with viral proteins to modulate SARS-CoV-2 replication.

## INTRODUCTION

The coronavirus disease 2019 (COVID-19) pandemic is caused by severe acute respiratory syndrome coronavirus-2 (SARS-CoV-2), which belongs to the genus *Betacoronavirus* in the *Coronavirinae* subfamily of the *Coronaviridae* family (1–3). The rapid transmission of COVID-19 has been a major global challenge. Similar to the other two β-category coronaviruses, SARS-CoV-2 harbors a positive-sense, single-stranded RNA genome of approximately 30 kb, with 80% and 50% homology to SARS-CoV and Middle East respiratory syndrome coronavirus (MERS-CoV), respectively (4).

Internal chemical modifications of viral RNA play key roles in the regulation of viral infection. N6-methyladenosine (m^6^A), 5-methylcytosine (m^5^C), and N4-acetylcytidine (ac4C) have been reported to be involved in the viral life cycle (5–10). m^6^A is one of the most abundant internal RNA modifications (11,12). The m^6^A machinery consists of “writers”, “erasers”, and “readers”. The writers, including methyltransferase-like (METTL) 3, METTL14, WT1-associated protein (WTAP), and other proteins, catalyze the transfer of the m^6^A modification (13–23). The “erasers” fat mass and obesity associated protein (FTO) and AlkB homolog 5 (ALKBH5) are m^6^A demethylases that remove the methyl groups from RNA (22–25). The “readers” contain a YTH motif that binds to m^6^A sites and play critical roles in mRNA stability (26–28), RNA processing (25), RNA structure (29), and translation (30,31).

The internal m^6^A modification of viral RNA was first identified in viruses that replicate in the cytoplasm, such as vesicular stomatitis virus, vaccinia virus, and reovirus, 40 years ago (32–36). However, the function of m^6^A began was only recent elucidated in hepatitis C virus (HCV), Zika virus (ZIKV), dengue virus, yellow fever virus, and West Nile virus (37,38). In viruses that replicate in the nucleus, such as human immunodeficiency virus (HIV), simian virus 40, Kaposi's sarcoma-associated herpesvirus, and influenza virus, viral m^6^A modifications have been shown to affect viral replication and gene expression (39–46). However, it is unclear whether SARS-CoV-2 contains m^6^A modifications although some potential m^6^A sites have been discovered by bioinformatics (47). At least 41 sites modified on the SARS-CoV-2 genome are potential sites of RNA modification, with frequent occurrence of the AAGAA motif, which is particularly enriched at genomic nucleotide positions 28,500–29,500 (48). However, although the m^5^C modification has been reported, the modification type is largely unknown (48).

Accordingly, in the current study, we investigated the presence of roles of m^6^A modification in SARS-CoV-2 RNA using methylated RNA immunoprecipitation sequencing (MeRIP-Seq). Overall, our findings demonstrated that the host m^6^A modification complex interacted with viral proteins and modulated SARS-CoV-2 replication.

## MATERIAL AND METHODS

### Virus, cell lines, and cell culture

SARS-CoV-2 was isolated from the bronchoalveolar lavage fluid of a patient (49) and passaged in monkey kidney cells (Vero-E6 cells) for eight generations. The titer of the SARS-CoV-2 working solution was 10^6^ PFU/mL, as determined by plaque assays in Vero-E6 cells. Vero-E6 cells (American Tissue Culture Collection [ATCC], Manassas, VA, USA; CRL-1586) and HEK293T cell (ATCC; CRL-11268) were cultured in Dulbecco’s modified Eagle’s medium 116 (Gibco, Gaithersburg, MD, USA) containing 10% fetal bovine serum (Gibco) at 37°C with 5% CO2.

### Plasmid construction and transfection

The RNA-dependent RNA polymerase (RdRp) plasmids pFlag-RdRp and pHA-RdRp were constructed by inserting the sequences of the RdRp open reading frame (ORF) into the vectors pXJ40-Flag and pXJ40-HA (Sigma-Aldrich, St. Louis, MO, USA), respectively. m^6^A methyltransferases and demethylase expression plasmids (pFlag-METTL3, pFlag-WTAP, pMETTL3, pMETTL14, and Flag-METTL3) were constructed by inserting the ORF sequences of the genes into the vectors pXJ40-Flag, pcDNA3.0, or pLenti-CMV-3XFlag. The plasmids HA-SUMO, HA-Ubi, HA-63, HA-48, and myc-ubc9 were kind gifts from Dr. Hanzhong Wang (Wuhan Institute of Virology, CAS). Plasmids were transfected into cells using Lipofectamine 2000 reagent (Invitrogen, Carlsbad, CA, USA; cat. no. 11668-019) according to the manufacturer’s instructions.

### Western blotting and antibodies

Cell lysates were prepared at the indicated times after transfection or infection and separated by gradient sodium dodecyl sulfate polyacrylamide gel electrophoresis (SDS-PAGE) on 10% gels, and proteins were then transferred to nitrocellulose membranes. The membranes were incubated with primary antibodies overnight at 4°C at the dilution suggested by the manufacturer’s protocols. The primary antibodies were as follows: anti-glyceraldehyde 3-phosphate dehydrogenase (GAPDH; cat. no. 60004-1-lg; Proteintech, Rosemont, IL, USA), mouse monoclonal anti-β-actin (cat. no. sc47778; Santa Cruz Biotechnology, Dallas, TX, USA), rabbit monoclonal anti-METTL3 (cat. no. 15073-1-AP; Proteintech), anti-METTL3 (cat. no. ab195352; Abcam, Cambridge, UK), anti-METTL14 (cat. no. SAB1104405; Sigma-Aldrich), anti-WTAP (cat. no. ab155655; Abcam), anti-ALKBH5 (cat. no. ab69325; Abcam), anti-FTO (cat. no. ab124892; Abcam), anti-Flag (cat. no. F1804-1 MG; Sigma-Aldrich), anti-HA (cat. no. H9658; Sigma-Aldrich), and mouse polyclonal anti-SARS-CoV-2 nonstructural protein (NP) (gift from Dr. Fei Deng, Wuhan Institute of Virology, CAS). The secondary antibodies including goat anti-mouse IgG and goat anti-rabbit IgG (AntiGene Biotech GmbH, Stuttgart, Germany) were incubated for 1 h at a dilution of 1:5000. Luminescent signals were detected using Tanon-5200 ChemiDoc MP imaging system (Tanon Science & Technology, Shanghai, China).

### Co-immunoprecipitation

Total proteins were collected at 48 h after transfection. Primary antibodies were mixed with supernatants of cell lysates (2 μg primary antibody per 1 mg protein sample) for 2 h at 4°C and then incubated with protein G agarose overnight at 4°C. Immunoprecipitated proteins were separated by SDS-PAGE on 12% gels and transferred nitrocellulose membranes, followed by incubation with primary and second antibodies. Protein detection was performed using Tanon 5200 ChemiDoc MP imaging system.

### Short hairpin RNA (shRNA) knockdown

shRNA knockdown was performed according to the protocol for shRNA-mediated gene silencing and Lentiviral Particle packaging from the Addgene website. Stable-knockdown Vero-E6 cell lines were screened using 10 μg/mL puromycin for selection. shRNA-specific primers were as follows: *METTL3* (shMETTL3-1: 5′-GCCAAGGAACAATCCATTGTT-3′, shMETTL3-2: 5′-CGTCAGTATATTGGGCAAGTT-3′), *FTO* (shFTO-1: 5′-TCACCAAGGAGACTGCTATTT-3′, shFTO-2: 5′-GATCCAAGGCAAAGATTTACT-3′).

### Immunofluorescence analysis

Immunofluorescence analysis were performed as previously described (50). Briefly, Vero-E6 cells were infected with SARS-CoV-2 (multiplicity of infection [MOI] = 0.01) and harvested 24 h postinfection. Cells were fixed in 4% paraformaldehyde overnight, permeabilized in 0.2% Triton X-100 for 10 min, washed three times with phosphate-buffered saline (PBS), and blocked in 3% bovine serum albumin for 1 h at room temperature. The cells were incubated with primary antibodies overnight at 4°C at the dilution suggested by the manufacturer’s protocol and stained with secondary antibodies (Alexa Fluor 488, Alexa Fluor 568) for 1 h after three washes with PBS. Nuclei were visualized with Hoechst 33258 at a dilution of 1:1000. The images were captured under a PerkinElmer VoX confocal microscope.

### Quantitative reverse transcription polymerase chain reaction (qRT-PCR)

Total RNA was extracted from SARS-CoV-2-infected Vero-E6 cells, and reverse transcription was performed using a HiSeript first-strand cDNA synthesis kit (Vazyme Biotech Co, Nanjing, China) according to the manufacturer’s instructions, followed by quantitative PCR with SYBR Green (Yeasen Biotech Co, Shanghai, China) on a CFX Connect Real-Time system (Bio-Rad Laboratories, Hercules, CA, USA).

### Formaldehyde-crosslinked RNA-immunoprecipitation (RIP)

RIP was conducted as previously described (51). Briefly, infected Vero-E6 cells were crosslinked by adding PBS containing 1% methanol-free formaldehyde and incubated for 10 min at 37°C. The reaction was terminated by adding 2.5 M glycine, and the cells were lysed with 400 μL RIP buffer (150 mM KCl, 25 mM Tris-HCl [pH 7.4], 5 mM ethylenediaminetetraacetic acid [EDTA], 0.5 mM dithiothreitol [DTT], 0.5% NP40, 100 U/mL RNase inhibitor, 100 uM phenylmethylsulfonyl fluoride [PMSF], and 1 μg/mL proteinase inhibitor) on ice for 10 min. The lysates were then centrifugated at 16,000 × *g* for 10 min, and supernatants were subjected to IP with IgG or anti-Flag antibodies overnight. Next, 30 μL protein-G agarose beads was added after washing three times with washing buffer (300 mM KCl, 25 mM Tris-HCl [pH 7.4], 5 mM EDTA, 0.5 mM DTT, 0.5% NP40, 100 U/mL RNase inhibitor, 100 μM PMSF, and 1 μg/mL proteinase inhibitor) and incubated with the indicated antibodies for 2 h at 4°C. RNA isolation was performed using TRIzol (Invitrogen, Carlsbad, CA, USA) for subsequent qualification.

### MeRIP-Seq

Total RNA was extracted from SARS-CoV-2-infected Vero-E6 cells and purified with an Oligo (dT) kit (Thermo Scientific, Wilmington, DE, USA). The polyA purified RNA was fragmented and subjected to IP using an m^6^A-specific antibody, followed by next-generation sequencing. MeRIP-Seq data were analyzed as described previously (44).

### MeRIP and northern blotting

For MeRIP and northern blotting, 400 μg total RNA from virus-infected Vero-E6 cells was incubated with an anti-m^6^A antibody (Synaptic Systems, Goettingen, Germany) or an IgG antibody in 300 μL IP buffer (150 mM NaCl, 0.1% NP-40, 10 mM Tris-HCl [pH 7.4]) for 2 h at 4°C. Thirty-five microliters magnetic beads (NEB; goat anti-rabbit magnetic beads; cat. no. S1432S) were added, and samples were washed three times and rotated 2 h at 4°C. The beads were washed six times and incubated with 300 μL elution buffer (5 mM Tris-HCl [pH 7.5], 1 mM EDTA [pH 8.0], 0.05% SDS, and 8.4 μL of 10 mg/mL proteinase K) for 1.5 h at 50°C. The RNA was purified using phenol/chloroform and precipitated with ethanol. For qRT-PCR, cDNA was synthesized using reverse transcriptase mix (Vazyme), and relative quantification was performed using specific primers. The data were normalized to the quantification cycle (Cq) values of *GAPDH*. For northern blotting, the purified RNA was run on a 1.0% agarose gel containing 2.2 M formaldehyde for 11 h at 35 V, followed by transfer to a Hybond-N+ membrane and ultraviolet crosslinking. Finally, membranes were hybridized with a DIG-labelled SARS-CoV-2 probe (nt 28,274–29,870), and probe detection was performed using a luminescence detection kit II (Roche) according to the manufacturer’s protocol. Signals were detected using a ChemiDoc MP imaging system (Tanon 5200).

### Sumoylation and ubiquitination assays

Sumoylation and ubiquitination assays were performed as previously described (52). Briefly, the indicated plasmids were cotransfected into HEK293T cells, and the cell lysates were harvested by centrifugation at 16,000 × *g* at 4°C for 10 min. Next, 50 μL protein G Dynabeads was incubated with 10 μg of the indicated antibodies for 2 h, followed by incubation with cell lysates overnight. The complexes were washed six times with PBS containing 0.02% Tween 20 and subjected to western blotting.

### Statistical analysis

The statistical analysis of qRT-PCR data was performed using two-tail unpaired t-tests in GraphPad Prism Software (GraphPad Software, La Jolla, CA, USA.). Data are presented as means ± standard deviations (n = 3). All experiments were repeated at least three times.

## RESULTS

### SARS-CoV-2 infection altered the expression patterns of m^6^A methyltransferases and demethylases

m^6^A methyltransferase and demethylases are mainly localized in the nucleus. Infection by viruses that replicate in the cytoplasm, such as EV71, HCV, and ZIKV, affects the expression and localization of methyltransferases and demethylase to facilitate their RNA m^6^A modifications, which promotes viral replication (37,38,51,53). To check whether SARS-CoV-2 infection had a similar effect, SARS-CoV-2-infected Vero-E6 cells were harvested. The expression of viral NP and m^6^A machinery proteins was assessed by western blotting with corresponding antibodies (Fig. 1A). Our results showed that the expression of METTL3 was increased at 48 h postinfection (hpi), whereas the expression levels of METTL14 and WTAP were not affected (Fig. 1A). The expression of the demethylase FTO decreased at 48 hpi, whereas that of ALKBH5 was not changed after infection. Moreover, the expression of the m^6^A binding proteins YTHDF1–3, YTHDC1, and YTHDC2 was not altered during SARS-CoV-2 infection (Fig.1A).

**Figure 1.**
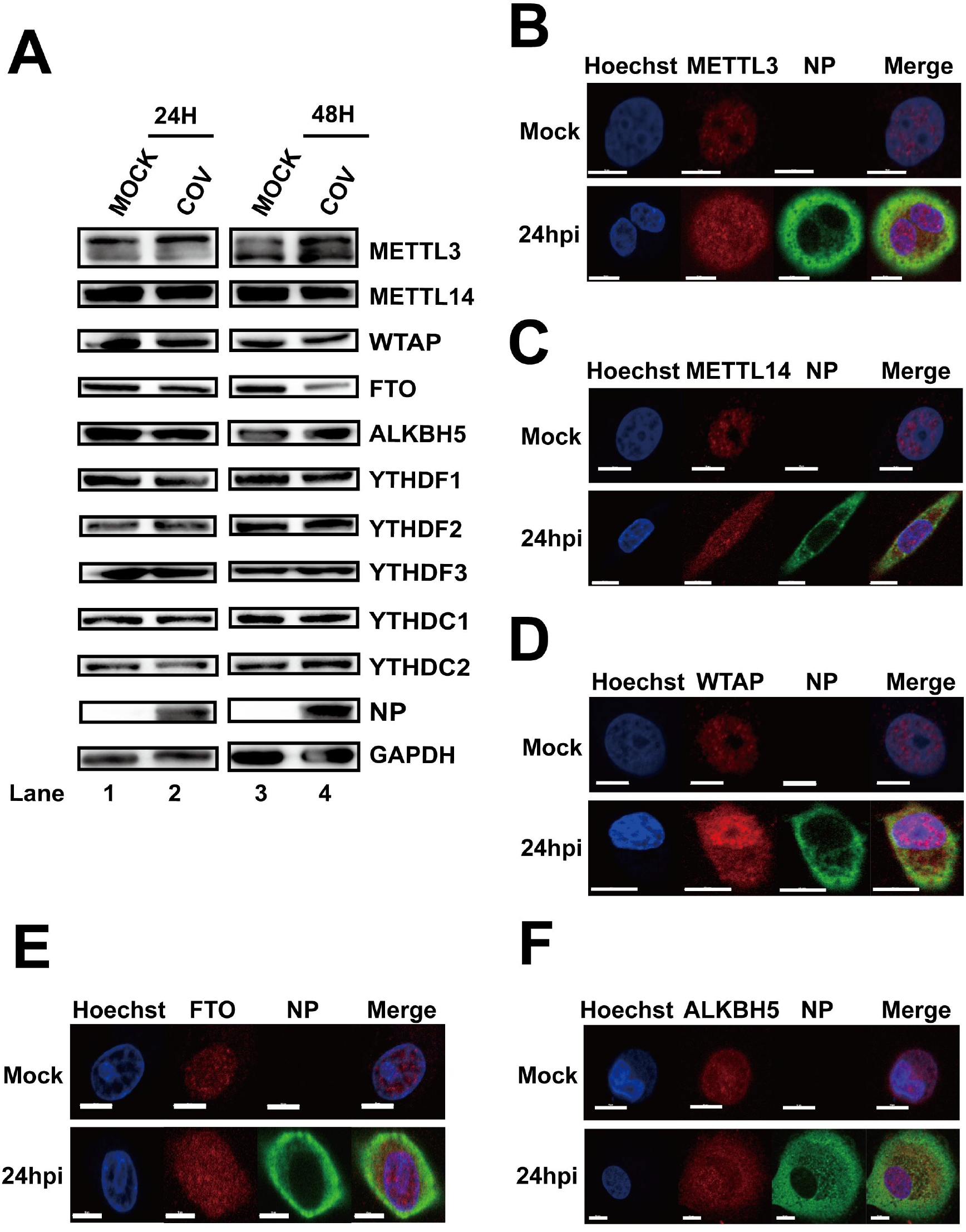
SARS-CoV-2 infection influenced the expression patterns of m^6^A-related proteins. Vero-E6 cells infected with SARS-CoV-2 (MOI = 0.01) were harvested at 24 and 48 hpi. Western blotting was performed with antibodies as indicated. GAPDH was used as a loading control. (B–F) Confocal microscopy images of SARS-CoV-2-or mock-infected Vero-E6 cells. The nucleus (blue) and virus protein NP (green) were labeled with Hoechst and anti-NP1-specific antibodies, respectively. The methyltransferases and demethylases were stained with antibodies as indicated. Scale bars, 5 μm.

Previous studies have shown that m^6^A methyltransferases and demethylases colocalize with nuclear speckle markers and that viral infection affects the subcellular localization of m^6^A-related proteins. Because SARS-CoV-2 infection affects the expression of METTL3 and FTO, we next determined the effects of SARS-CoV-2 infection on the localization of methyltransferases and demethylases. Consistent with previous results, methyltransferases and demethylases were detected mostly in the nucleus under normal conditions (Fig. 1B–F). However, METTL3, METTL14, WTAP, ALKBH5, and FTO were all present in both the nucleus and cytoplasm after infection (Fig. 1B–F). The colocalization of methyltransferases and demethylases with viral protein NP implied that these proteins may interact with SARS-CoV-2 RNA in the cytoplasm. The above results provided evidence that SARS-CoV-2 may be modified by the host m^6^A machinery.

### SARS-CoV-2 RNA contained m^6^A modifications

At least 41 sites on the SARS-CoV-2 genome contain potential sites of RNA modifications, harboring the AAGAA motif (48). However, the modification type is largely unknown. To investigate whether SARS-CoV-2 RNA was m^6^A modified, total RNAs were purified from large-scale batches of SARS-CoV-2-infected Vero-E6 cells, and MeRIP was then performed with m^6^A-specific antibodies. The MeRIP RNA was subjected to northern blotting with SARS-CoV-2 probes spanning nt 28,274–29,870. SARS-CoV-2 RNA was then pulled down using anti-m^6^A antibodies (Fig. 2A), indicating that SARS-CoV-2 contained m^6^A residues. To further confirm the above results and map the m^6^A modification status in the SARS-CoV-2 RNA genome, MeRIP-Seq was performed. Five m^6^A peaks were identified in the 5′ end (nt 36–753 and nt 1023–1324) and the 3′ end (nt 27,493–27,913, nt 28,475–28,706, and nt 28,944–29,751; Fig. 2B–D), which were located in the ORF1ab-, N-, and ORF10-coding regions (Fig. 2E). These results implied that SARS-CoV-2 RNA was marked by m^6^A modification during infection.

**Figure 2.**
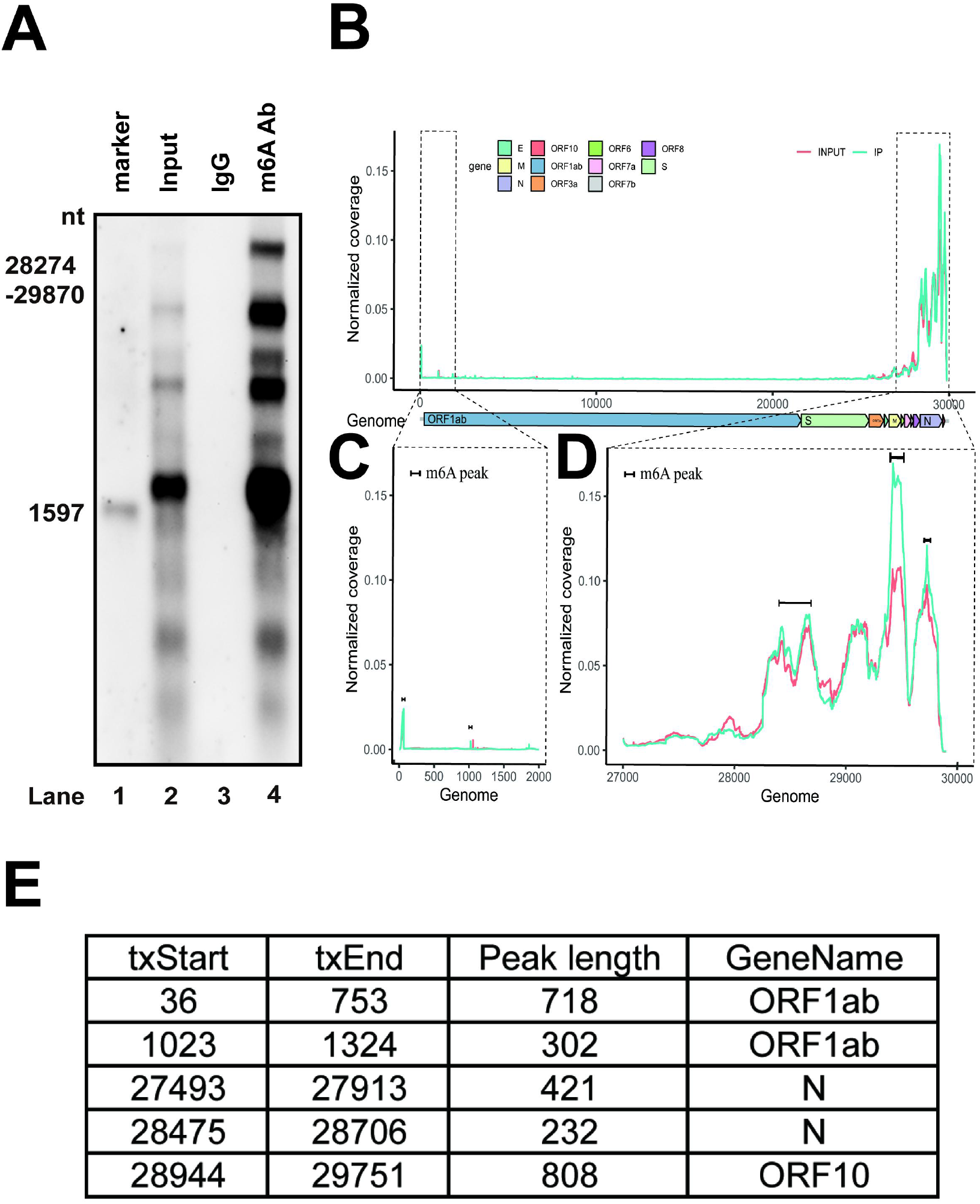
SARS-CoV-2 genomic RNA harbored m^6^A modifications. (A) MeRIP and northern blotting. RNAs from virus-infected Vero-E6 cells were incubated with IgG or anti-m^6^A-specific antibodies as indicated. Immunoprecipitated RNAs were resolved on 1% agarose gels containing 2.2 M formaldehyde and transferred to Hybond-N^+^ membranes, followed by RNA signal detection with SARS-CoV-2-specific probes spanning from nt 28,274 to nt 29,870. (B–D) MeRIP-Seq. Fragmented total RNAs from SARS-CoV-2-infected Vero-E6 cells were subjected to IP with anti-m^6^A-specific antibodies, followed by next-generation sequencing. Methylation coverage of the full-length SARS-CoV-2 RNA is shown. Representative of n = 3 determinations. (E) Table of m^6^A peak regions. The five peak regions are shown.

### METTL3 regulated m^6^A modification of SARS-CoV-2 RNA and virus replication

The host methyltransferases and demethylases are involved in the m^6^A modification of EV71, HCV, ZIKV, and HIV because these viruses do not encode any enzymes with m^6^A methyltransferase activity (37-41,51,54). To determine whether host m^6^A machinery was responsible for the viral m^6^A modification, the FLAG-tagged *METTL3* gene was expressed in Vero-E6 cells by transfection (Fig. 3A). qRT-PCR of the *RdRp* gene was performed following formaldehyde-crosslinked RIP using an anti-FLAG antibody to pull down METTL3-bound RNAs. SARS-CoV-2 RNAs were pulled down by METTL3 (Fig. 3B), indicating that SARS-CoV-2 RNA could interact with METTL3. We next knocked down endogenous *METTL3* in Vero-E6 cells using shRNA (Fig. 3D). m^6^A abundance in SARS-CoV-2 RNA was detected by using qRT-PCR (Fig. 3E) or northern blotting (Fig. 3F) after MeRIP. We found that silencing METTL3 by shRNA resulted in decreased abundance of m^6^A in SARS-CoV-2 RNA (Fig. 3D and E). In contrast, overexpression of METTL3 by transfection increased the abundance of m^6^A-bound SAR-CoV-2 RNAs (Fig. 3C). To further confirm our results, MeRIP-Seq was performed after *METTL3* knockdown (Fig. 3G and Fig. S4). Our results showed that the methylation peaks were not changed, but that the frequency of methylation was significantly decreased, suggesting that the m^6^A modification levels in the SARS-CoV-2 genome were linked to METTL3 expression. Taken together, these results indicated that METTL3 acted as a methyltransferase in the viral genome.

**Figure 3.**
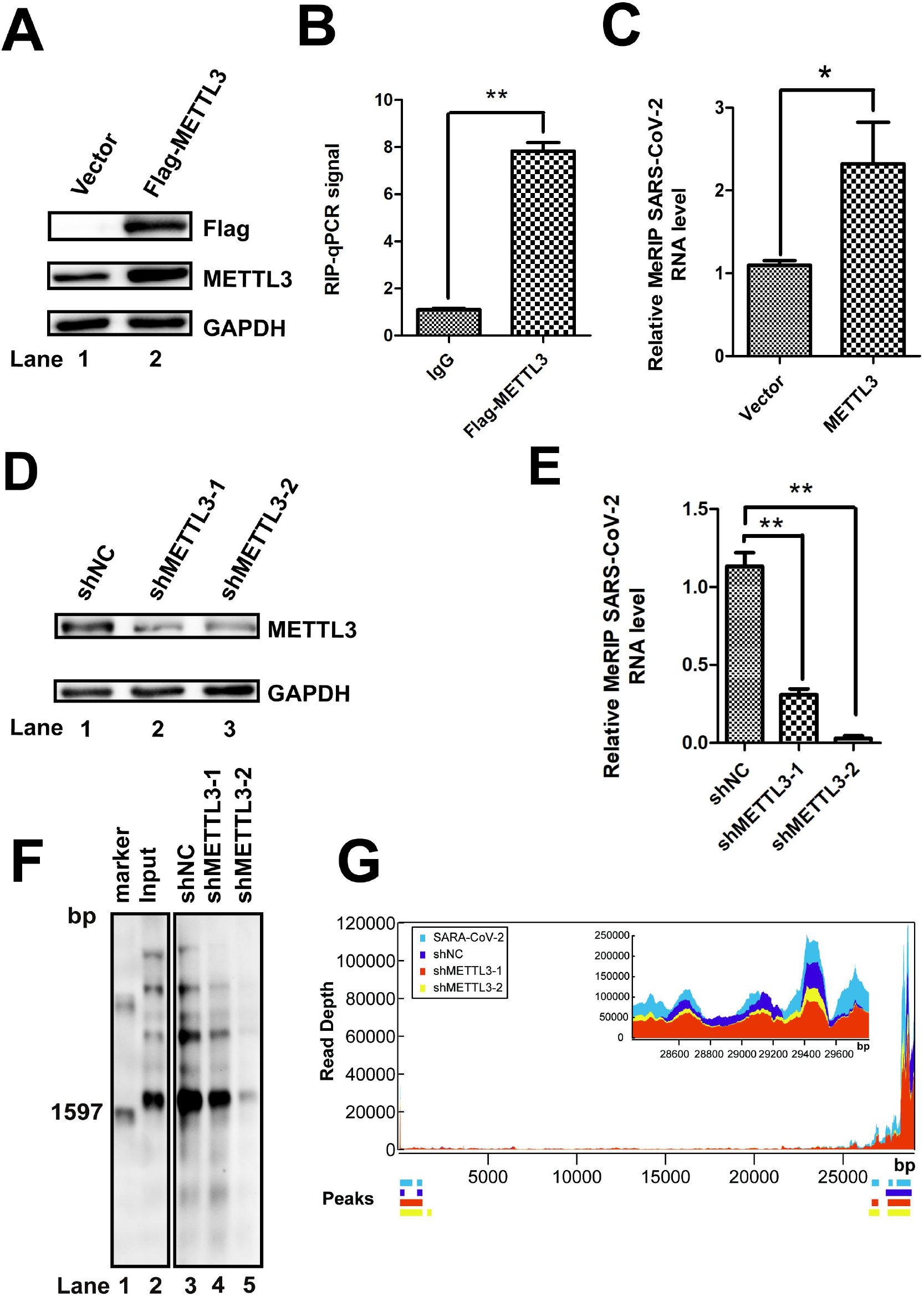
METTL3 catalyzed the m^6^A modification of SARS-CoV-2. (A and D) Western blotting. METTL3 was knocked down by shRNA (D) or overexpressed (A) in Vero-E6 cells. The expression of METTL3 was checked using anti-METTL3 (A and D) or anti-Flag antibodies (A), as indicated. Vector-transfected cells were used as a control. (B) Formaldehyde-RIP qRT-PCR. Cell lysates from formaldehyde-crosslinking were subjected to IP with IgG or anti-Flag antibodies. qRT-PCR was performed to quantify SARS-CoV-2 RNA. IgG was used as a negative control. Unpaired Student’s *t*-tests were performed, and data are presented as means ± standard errors of the means (n = 3). \*\**P* ≤ 0.01. (C and E) MeRIP-qPCR. RNA was extracted from SARS-CoV-2-infected Vero-E6 cells in which *METTL3* was overexpressed (C) or knocked down by shRNA (E). Me-RIP was performed, and SARS-CoV-2 RNA was quantified by qRT-PCR. Unpaired Student’s *t*-tests were performed, and data are presented as means ± standard errors of the means (n = 3). \*\**P* ≤ 0.01. (F) MeRIP and northern blotting. RNAs were harvested from SARS-CoV-2-infected Vero-E6 cells in which METTL3 was knocked down by shRNA. MeRIP and northern blotting were performed. (G) MeRIP-Seq. Total RNA was isolated from SARS-CoV-2-infected Vero E6 cells in which METTL3 was knocked down.

Viral protein expression and progeny virus production by HIV, HCV, and EV71 are affected by the expression of endogenous methyltransferases or demethylases (37,39-41,51). Endogenous *METTL3* (Fig. 4A) or FTO (Fig. 4B) in Vero-E6 cells was knocked down by specific shRNAs, followed by SARS-CoV-2 infection to check whether METTL3 or FTO affected viral replication. Viral RNA copy numbers were quantified by qRT-PCR of the *N* or *RdRp* gene using standard protocols. We found that efficient knockdown of *METTL3* resulted in not only significantly decreases in viral *N* and *RdRp* gene copy numbers (Fig. 4C and D) but also decreased expression of NP (Fig. 4A). However, knockdown of *FTO* had the opposite effect (Fig. 4B, E, F). These results suggested that the m^6^A methyltransferase METTL3 was linked to efficient SARS-CoV-2 replication.

**Figure 4.**
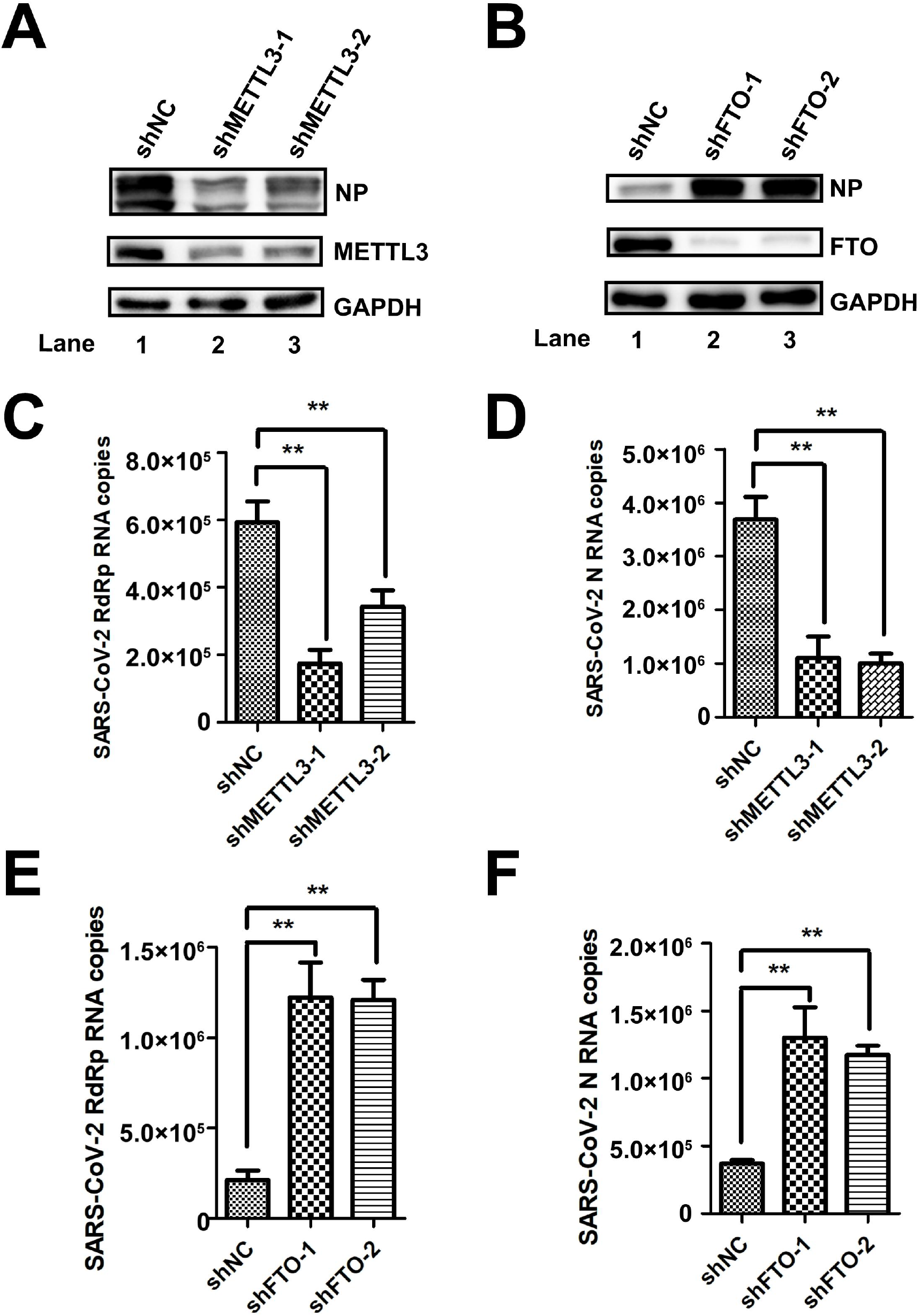
METTL3 regulated SARS-CoV-2 replication. (A and B) Western blotting. *METTL3* and *FTO* was knocked down in Vero-E6 cells by shRNA. The expression of METTL3 protein and viral NP was detected by western blotting with specific antibodies. (C–F) qRT-PCR. Total RNA was isolated from SARS-CoV-2-infected Vero-E6 cells in which METTL3 and *FTO* was knocked down by shRNA as indicated. SARS-CoV-2 RNA was quantified using qRT-PCR with specific primers targeting *N* and *RdRp* genes. *GAPDH* was used as a control. Unpaired Student’s *t*-tests were performed. Data are presented as means ± SEMs (n = 3). \**P* ≤ 0.05.

### SRAS-CoV-2 RdRp interacted with METTL3 and influenced its expression

In our previous study, METTL3 modulated EV71 replication by interacting with EV71 polymerase 3D and regulating 3D sumoylation and ubiquitination (51). To investigate whether there was a similar mechanism in SARS-CoV-2, pFlag-METTL3 and pHA-RdRp were cotransfected into HEK293T cells. The IP experiment with anti-Flag antibodies, followed by staining with anti-HA or vice versa, showed that METTL3 interacted with SARS-CoV-2 RdRp protein in the absence or presence of RNase A (Fig. 5A and B, Fig. S1A and B). In addition, our study showed that RdRp interacted with the methyltransferase complex (Fig. S2). To further confirm our results, the colocalization of METTL3 and RdRp was checked after cotransfection of the cells with the two plasmids. Notably, METTL3 was distributed in both the nucleus and cytoplasm when RdRp was co-expressed (Fig. 5E).The colocalization of METTL3 and RdRp supported the interaction between these two proteins, which bound to viral RNAs. Moreover, METTL3 expression was increased as more RdRp was expressed by transfection or vice versa (Fig. 5C and D), suggesting that the abundance of METTL3 and RdRp influenced the expression of RdRp or METTL3.

**Figure 5.**
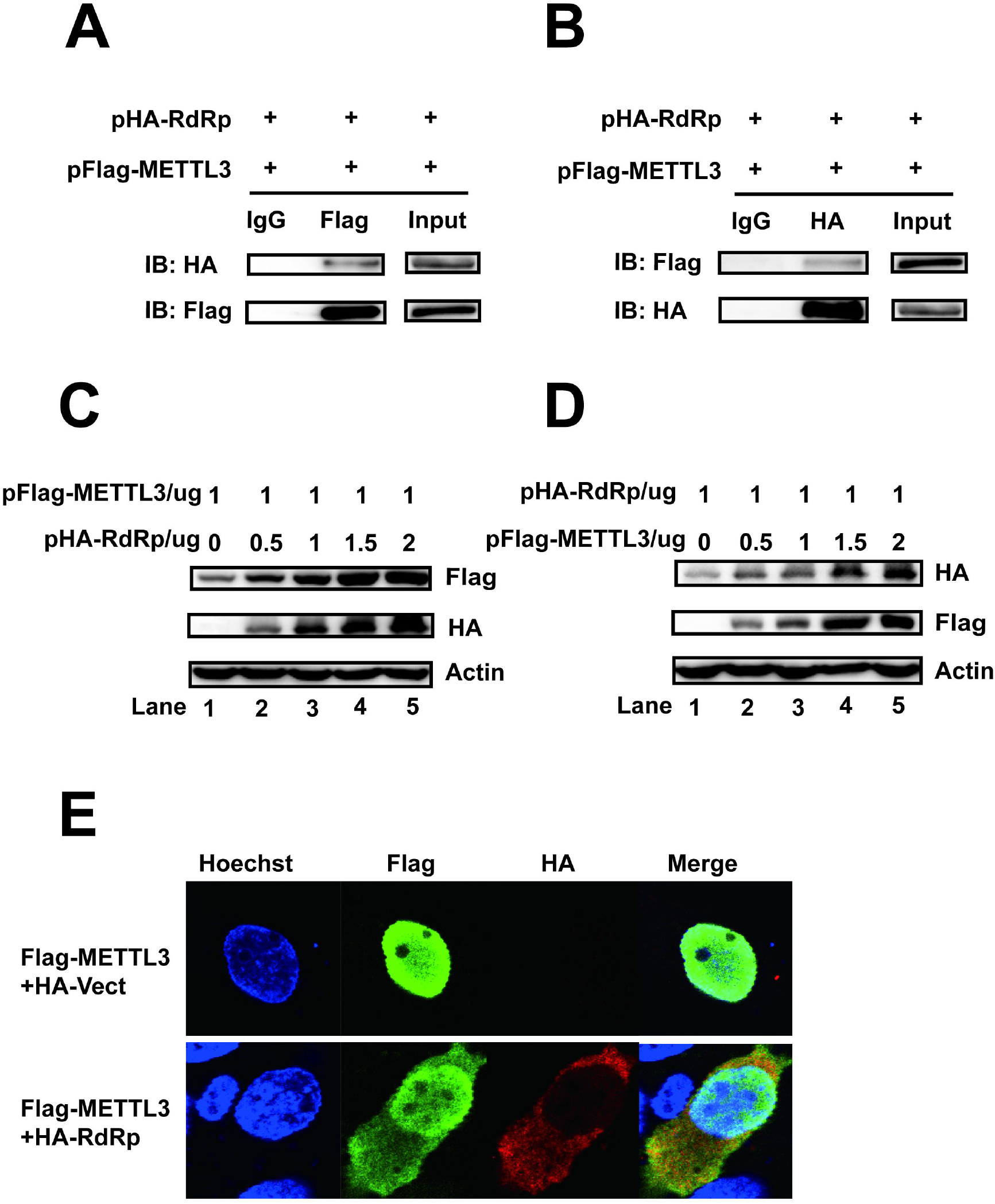
METTL3 interacted with SARS-CoV-2 RdRp. (A and B) Western blotting. pFlag-METTL3 and pHA-RdRp were cotransfected into HEK293T cells, and co-IP was performed with anti-HA (A) or anti-Flag (B) antibodies. IgG antibodies were used as a control. The IP samples were pulled down with anti-Flag (A) or anti-METTL3 (B) antibodies. (C and D) Western blotting. HEK293T cells were transfected with 1 μg pFlag-METTL3 (C) or pHA-RdRp (D) together with different amounts of pHA-RdRp (C) or pFlag-METTL3 (D) (0, 0.5, 1, 1.5, and 2 μg, respectively) in six-well plates. The expression of METTL3 and RdRp was detected by western blotting. (E) Confocal microscopy images. HEK293T cells were transfected with pFlag-METTL3 with or without HA-RdRp transfection. Costaining was performed using anti-Flag (green) and anti-HA antibodies (red). The nucleus (blue) was stained with Hoechst.

### SRAS-CoV-2 RdRp regulated METTL3 sumoylation and ubiquitination

To analyze how viral protein RdRp affected the m^6^A machinery components, we first checked the RNA abundance of all the m^6^A writers, erasers, and readers after different infection times, as indicated (Fig. 6A). The RNA abundances of *METTL3*, *METT14*, *WTAP*, *ALKBH5*, *FTO*, and *YTH* were not changed (Fig. 6A and Fig. S3), indicating that SARS-CoV-2 did not influence the RNA levels of m^6^A-related proteins. Post-transcriptional modifications, such as ubiquitination and sumoylation, affect METTL3 protein abundance and function. EV71 3D protein interacted with METTL3 and affected the expression and localization METTL3, similar to the results observed for SARS-CoV-2 infection. We next investigated whether RdRp affected the modification of METTL3. To this end, METTL3, pFlag-RdRp, HA-SUMO-1, and myc-Ubc-9 were transfected into HEK293T cells. The western blotting results showed that sumoylation of METTL3 was reduced in the presence of RdRp expression (Fig. 6B). Cotransfection with pMETTL3, Flag-RdRp, and HA-Ub resulted in decreased ubiquitination of METTL3 (Fig. 6C). Further experiments showed that overexpression of RdRp resulted in decreased K48-linked ubiquitination (Fig. 6D) and K63-linked ubiquitination (Fig. 6E), which could explain how RdRp regulated METTL3 expression.

**Figure 6.**
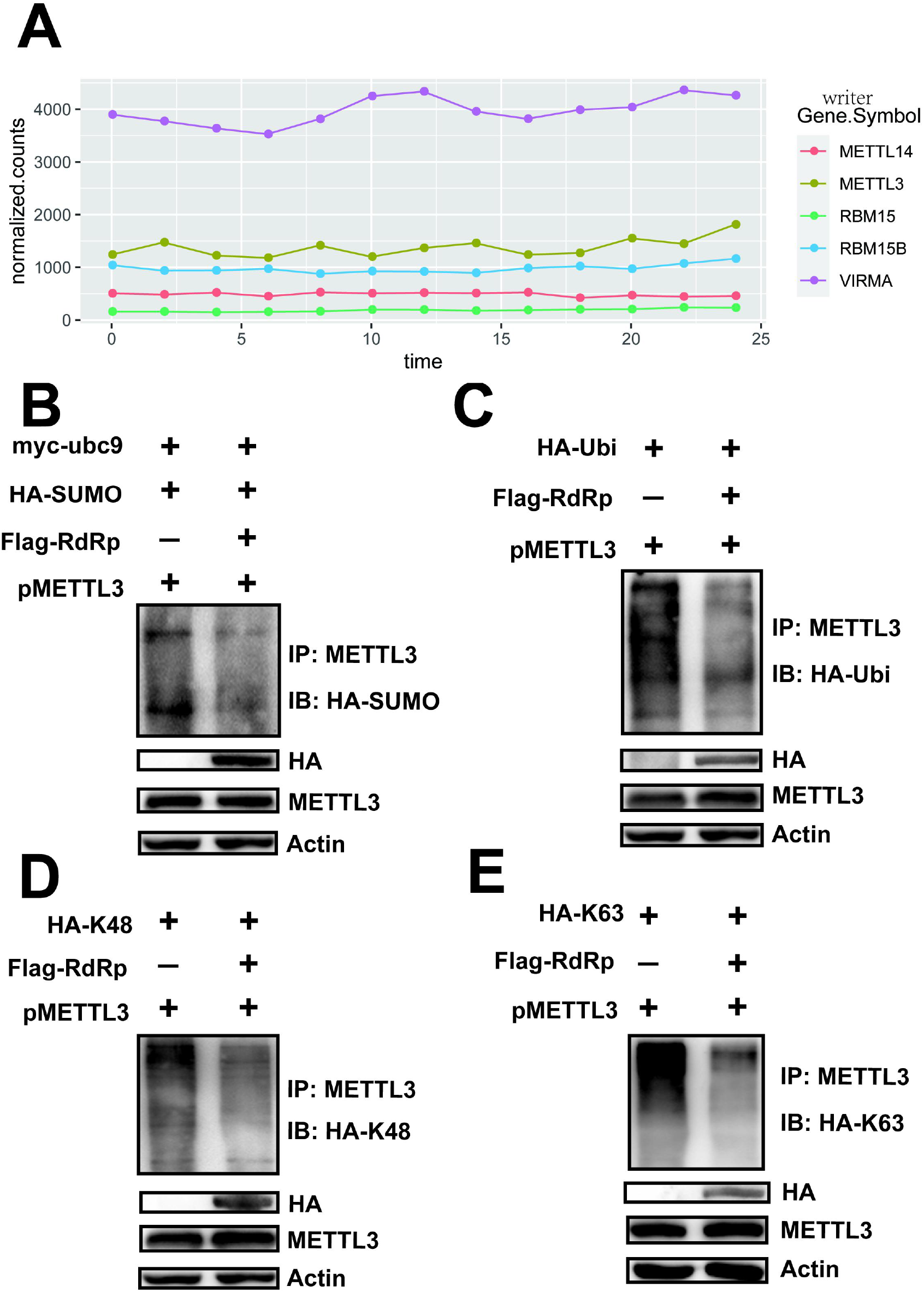
RdRp expression regulated the sumoylation and ubiquitination of METTL3. (A) RNA expression of host methyltransferases. Total RNA was harvested from SARS-CoV-2-infected Vero-E6 cells every 2 h, as indicated. The mRNAs were separated and subjected to next-generation sequencing. RNA levels of host methyltransferases were normalized according to the sequencing reads. (B) Sumoylation assay. *METTL3* was overexpressed in HEK293T cells by transfection with pMETTL3, followed by transfection with pFlag-RdRp, pHA-SUMO-1, and pMyc-Ubc9. IP and immunoblot analyses were performed using the indicated antibodies for the sumoylation assay. (C–E) Ubiquitination assay. HEK293T cells were transfected with pFlag-RdRp, pHA-Ubi, pHA-K48, and pHA-K63 after *METTL3* overexpression. IP and immunoblot analyses were performed using the indicated antibodies.

## DISCUSSION

In the current study, we demonstrated that SARS-CoV-2 RNA underwent m^6^A modification by host m^6^A machinery. The expression and localization of host m^6^A components were altered during SARS-CoV-2 infection. Knockdown of METTL3 decreased the replication of SARS-CoV-2, indicating that m^6^A modification played key roles in viral replication. Further studies showed that the viral polymerase RdRp interacted with METTL3 and regulated its sumoylation and ubiquitination to affect its expression and localization. Overall, our study showed that SARS-CoV-2 RNA was m^6^A modified and that METTL3 played a role in regulating viral replication.

RNA modification, such as m^6^A, m^5^C, and ac4C, regulates viral protein expression and progeny virus production (5–10). m^6^A modification has been identified in RNA viruses replicating both in the nucleus and cytoplasm (5,9,55). In our previous study, we found that coronavirus WIV-1 harbored an m^6^A modification (data not shown). Therefore, we assessed the internal m^6^A modification status of SARS-CoV-2 RNA by MeRIP-Seq; the results demonstrated that m^6^A peaks were mainly distributed in both 5′ and 3′ ends spanning the ORF1ab-, N-, and ORF10-encoding regions. The m^6^A modification pattern of SARS-CoV-2 is very similar to that of host mRNAs (12,56), but different from that of EV71 virus, whose m^6^A sites are distributed in coding regions of the middle of viral genome (51). The m^6^A motif (RRACH) is different from reported RNA modification sites harboring an AAGAA motif. Notably, 44 m^6^A motifs were found in the five enriched peaks, most of which were distributed in the N gene region.

SARS-CoV-2 infection resulted in not only elevated expression of METTL3 but also altered distribution in both the nucleus and cytoplasm. We also found that METTL14, WTAP, ALKBH5, and FTO colocalized with the viral protein NP, supporting that SARS-CoV-2 infection affected the m^6^A methyltransferase and demethylases. The colocalization of viral NP and host m^6^A proteins supported that the m^6^A modification machinery could modify cytoplasmic SARS-CoV-2 RNA during infection. In our previous study, we found that METTL3 interacted with EV71 polymerase 3D protein (51).

Cotransfection of cells with METTL3 and 3D resulted in both nuclear and cytoplasmic distribution of METTL3, implying that 3D played roles in the distribution of METTL3 in the cytoplasm; however, the specific mechanism is still unknown. In the current study, we found that the SARS-CoV-2 RdRp protein induced the expression and cytoplasmic distribution of METTL3.

Most RNA viruses that replicate in the cytoplasm, including ZIKV, West Nile virus, and EV71, hijack the host m^6^A machinery to modify the RNA and therefore do not encode methyltransferase (37,38,51). SARS-CoV-2 nonstructural proteins NSP14 and NSP16 have methyltransferase function and play key roles in the m^7^G cap and 2’-O-methylation modification (57–59). Our study showed that METTL3 interacted with SARS-CoV-2 RNA. Notably, the expression of METTL3 is linked to the m^6^A modification level of SARS-CoV-2 RNA. Knocking down METTL3 resulted in decreased m^6^A modification of SARS-CoV-2 RNA, which was detected either by MeRIP northern blotting or by MERIP-Seq. However, overexpression of METTL3 resulted in elevated m^6^A modification, suggesting that METTL3 may be the methyltransferase modifying viral RNA.

METTL3 is a multifunctional protein that functions during EV71 infection. Viral RpRd 3D protein binds to the methyltransferase complex, and METTL3 regulates the ubiquitination of 3D to promote viral replication (51). In this study, knockdown of METTL3 resulted in decreased SARS-CoV-2 replication; this result could be explained by the absence of METTL3 methyltransferase activity, which catalyzes the methylation of viral RNA. To address whether key proteins of SARS-CoV-2 interacted with m^6^A components to facilitate virus replication, we checked the interactions of METTL3 with viral RdRp, which bound to viral RNA. The results showed that METTL3 not only interacted with RdRp but also promoted RdRp expression; the opposite result was also true. In contrast to EV71 3D protein, for which post-translational modification was modulated by METTL3, SARS-CoV-2 RdRp expression altered the localization pattern of METTL3. The distribution of METTL3 in the presence of RdRp expression confirmed the interaction between METTL3 and RdRp and may explain the presence of METTL3 in the cytoplasm during SARS-CoV-2 infection. Sumoylation and ubiquitination affect the function and expression of METTL3, respectively (60). To elucidate how RdRp expression increased the expression of METTL3, we checked the protein modification of METTL3. The results showed that RdRp expression decreased the sumoylation and overall ubiquitination levels. Moreover, K48- and K63-linked ubiquitination levels were reduced. These data supported that RdRp not only promoted methyltransferase activity but also increased METTL3 expression by decreasing its ubiquitination.

In summary, our results provided evidence that the host m^6^A machinery interacted with viral key proteins to facilitate the replication of SARS-CoV-2. First, METTL3 functioned as a methyltransferase, adding the m^6^A modification to viral RNA. Second, METTL3 interacted with viral RdRp, which resulted in METTL3 distribution both in the nucleus and cytoplasm. Importantly, RdRp boosted the expression of METTL3 by altering the ubiquitination pattern through an unknown mechanism (Fig. 7). Further studies are required to elucidate this mechanism. The functional m^6^A sites on the SARS-CoV-2 RNA were not defined in our study yet, which is under investigation.

**Figure 7.**
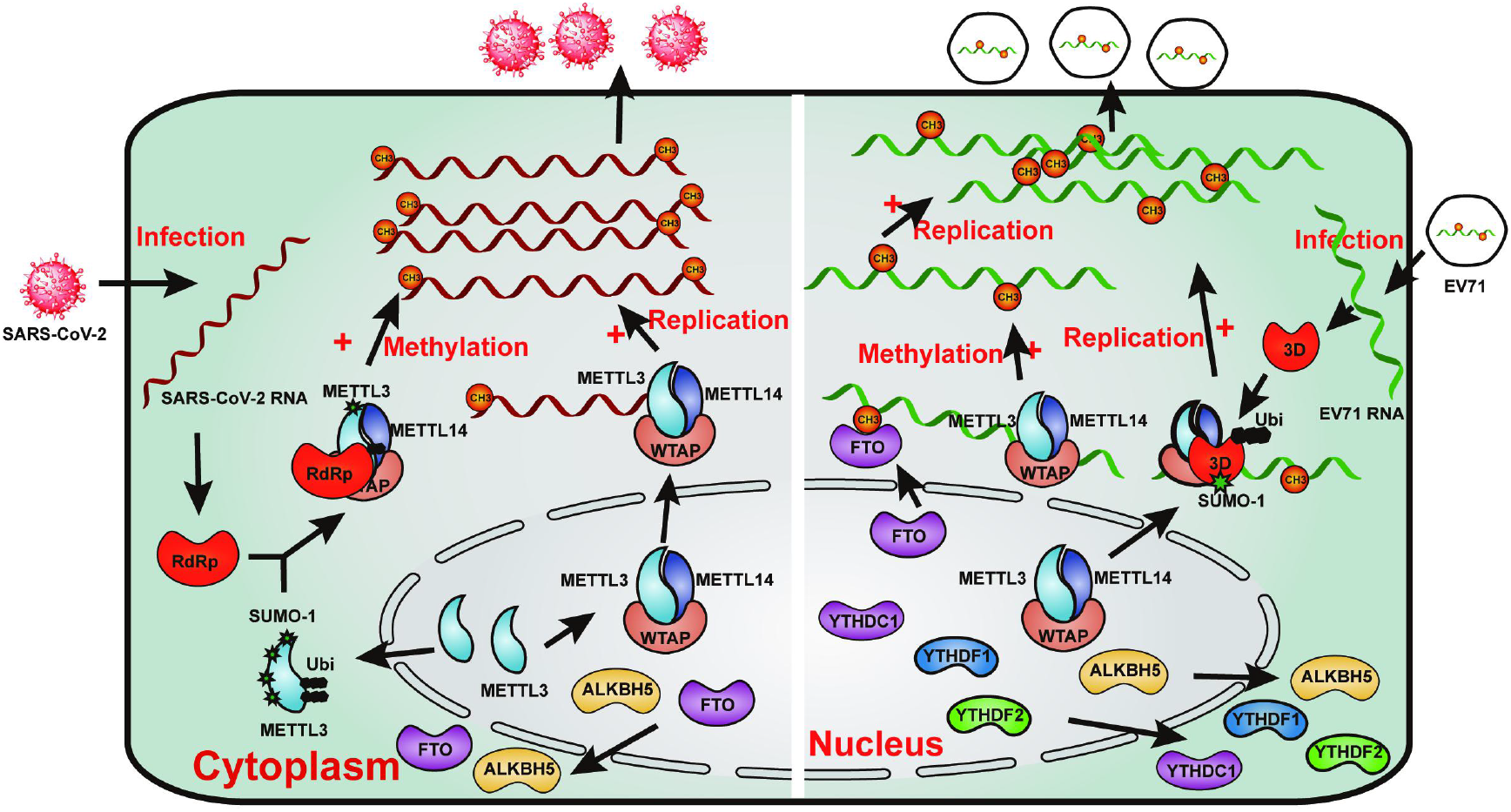
Model showing how the host m^6^A machinery modulates SARS-CoV-2 and EV71 replication.

## DATA AVAILABILITY

SARS-CoV-2 sequence data that support the findings of this study have been deposited in GISAID (https://www.gisaid.org/) with accession numbers EPI_ISL_402124, EPI_ISL_402127– EPI_ISL_402130, and EPI_ISL_402131; GenBank with accession numbers MN996527–MN996532; and National Genomics Data Center, Beijing Institute of Genomics, Chinese Academy of Sciences (https://bigd.big.ac.cn/databases?lang=en) with accession numbers SAMC133236–SAMC133240 and SAMC133252.

## ACKNOWLEDGEMENT

We are grateful to Lei Zhang and Ding Gao of the Core Facility and Technical Support at the Wuhan Institute of Virology, CAS for their help with next-generation sequencing and confocal microscopy.

## FUNDING

This work was supported by the Ministry of Science and Technology (grant nos. 2020YFC0845800 and 2020YFC0842500 to WG), Chinese Academy of Sciences (grant no. 2020YJFK0107 to WG), and Chinese Academy of Engineering (grant no. 2020-ZD-15 to WG). The funders had no role in the design, interpretation, or submission of this work for publication. Funding for open access charge: Chinese Academy of Sciences (grant no. 2020YJFK0107).

## CONFLICT OF INTEREST

None declared.

## Notes

https://www.gisaid.org/

https://bigd.big.ac.cn/databases?lang=en

### Competing Interest Statement

The authors have declared no competing interest.

