## Supplementary material for "Methyltransferase-like 3 modulates severe acute respiratory syndrome coronavirus-2 RNA N6-methyladenosine modification and replication": Fig S1,Fig S2,Fig S3,Fig S4

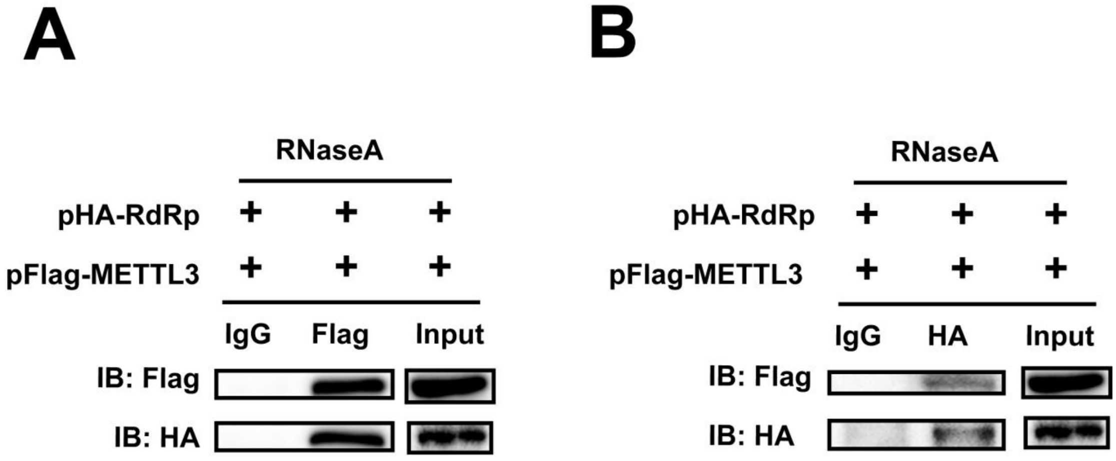


**Figure S1**. METTL3 interacted with SARS-CoV-2 RdRp in a RNA-independent way. (**A** & **B**) Western blotting. pFlag-METTL3 and pHA-RdRp were co-transfected into HEK293T cells and cell extracts were digested with RNaseA for 15min at 37℃. Co-IP was performed with anti-Flag (A) or anti-HA (B) antibodies. IgG antibody was used as a control. The immunoblots were visualized by the indicated antibodies.

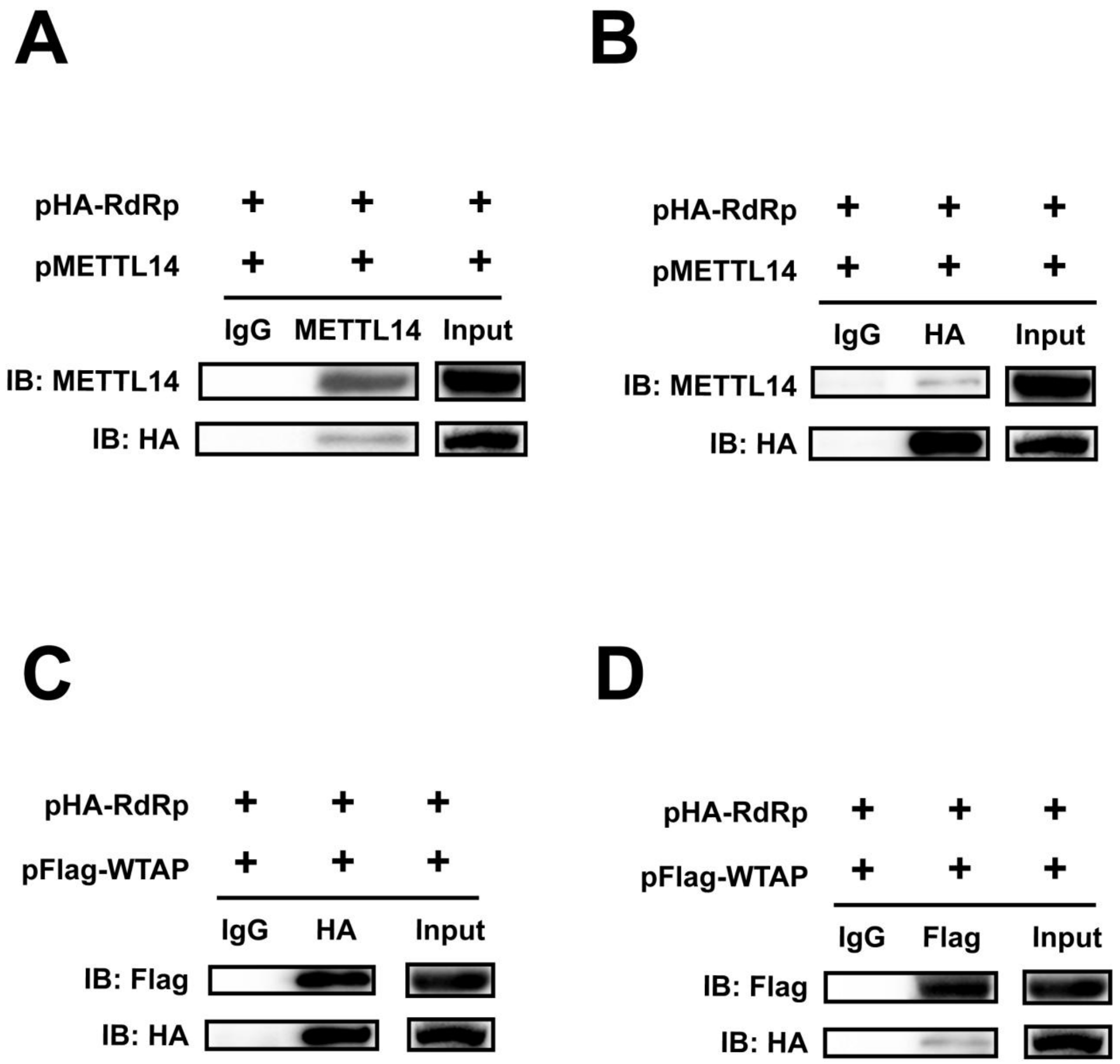


**Figure S2**. SARS-CoV-2 RdRp interacted with the methyltransferase complex.(A-D)Western blotting. HEK293T cells were co-transfected with pMETTL14 and pHA-RdRp(A & B) or pFlag-WTAP and pHA-RdRp(C & D).Co-IP was performed with anti-METTL14 (A) or anti-HA (B & C) or anti-Flag (D) antibodies. IgG antibody was used as a control. The immunoblots were visualized by the indicated antibodies.

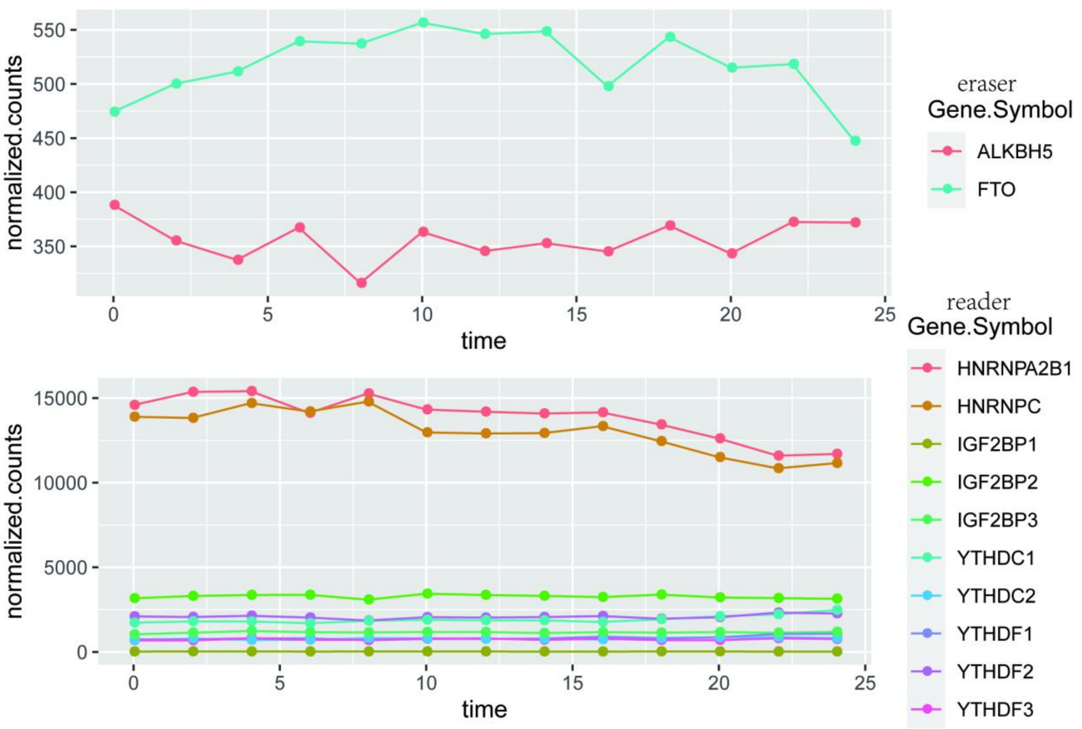


**Figure S3**. RNA expression of host demethylases and m6A binding protiens. Total RNA were harvested from SARS-CoV-2 infected Vero E6 cells every 2h as indicated. The mRNAs were separated and subjected to next generation sequencing. The RNA level of host demethylases and m6A binding protiens were normalized according to the counts.

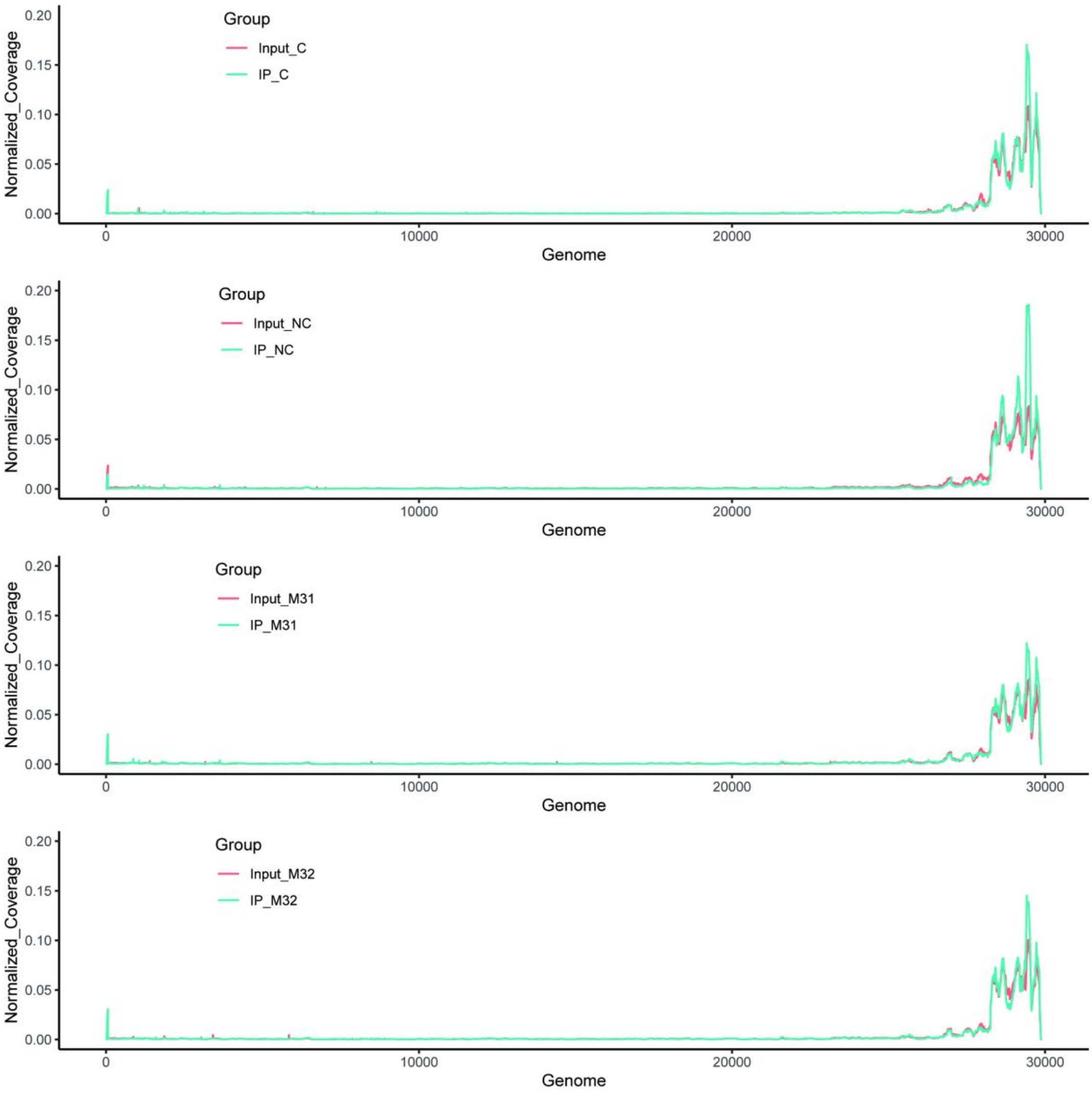


**Figure S4**. Effect of METTL3 gene knockdown in m6A methylation levels. Total RNAs were isolated from SARS-CoV-2 infected Vero E6 cells in which METTL3 was knocked down,then subjected to IP with m6A-specific antibody, followed by next-generation sequencing. In order to make the samples comparable, IP reads mapping to virus genome were normalized to the total number of sequenced reads. Peaks represent m6A-enriched regions detected by MACS2.
